# Learning Fragmentation Physics or Exploiting Sequence Priors? Benchmarking Bias in Deep Learning Models for De Novo Peptide Sequencing

**DOI:** 10.64898/2026.06.23.734131

**Authors:** Jiayi Li, Hannes Röst

## Abstract

Deep learning models have advanced de novo peptide sequencing, but their predictions may reflect both physics-based spectral evidence and learned peptide-sequence priors. Systematically measuring such prior-associated behavior is important for benchmarking model robustness beyond conventional proteomics data. Here, we introduce the Prior Bias Index (PBI), a general framework for measuring the extent to which model behavior shifts toward prior-associated reference patterns under controlled conditions, and implement it as DeNovo-PBI, a benchmark for quantifying prior bias in de novo peptide sequencing models. DeNovo-PBI combines benchmark dataset construction, *in silico* sequence and spectral perturbation workflows, PBI-based metrics, and analysis algorithms to evaluate three forms of prior-associated behavior: sequence-distribution dependence, database amino-acid-pair order preference, and mutation-group prediction consistency under shared sequence context. In addition to experimentally acquired peptide spectra, we generated *in silico* spectra from random, natural, and mutated peptide sequences and selectively removed fragment ions that distinguish N-terminal residue orders. Across these assays, deep learning models showed peptide-sequence-distribution-dependent performance and strong directional amino-acid-pair order preferences even when order-diagnostic spectral evidence was removed. DeNovo-PBI provides a quantitative benchmark for measuring, comparing, and interpreting learned bias in de novo peptide sequencing models.

## Introduction

Mass spectrometry has become a central technology for large-scale identification and quantification of proteins in complex biological samples, where peptide sequences are inferred from tandem mass spectrometry (MS/MS) spectra generated by peptide fragmentation^1,2^. In conventional workflows, peptide identification is commonly performed by matching spectra to known protein sequence databases^3,4^. However, database search can miss peptides that are not represented in the searched database, including those arising from unknown proteins, sequence variation, or unexpected modifications. De novo peptide sequencing provides a complementary strategy by directly inferring peptide sequences from MS/MS spectra without requiring a predefined database, making it particularly valuable for analyzing peptides outside the searched sequence space^5,6^.

Early de novo peptide sequencing methods primarily relied on heuristic search, graph-based algorithms, or dynamic programming to assemble peptide sequences from MS/MS spectra, as exemplified by tools such as Lutefisk^5^, SHERENGA^7^, and PEAKS^8^. Novor^9^ further improved de novo sequencing by combining dynamic programming with machine-learning-assisted scoring of peptide-spectrum evidence. More recently, deep learning approaches have reframed de novo sequencing as a learned prediction problem. Models such as DeepNovo^10^, SMSNet^11^, Casanovo^6,12^, and InstaNovo^13^ use neural network models to map MS/MS spectra to peptide sequences, achieving increasingly strong peptide- and amino-acid-level accuracy. Accordingly, existing benchmarks have largely focused on evaluating sequencing accuracy across datasets, species, instruments, and model architectures, establishing the performance gains of modern de novo sequencing methods^10,14^.

However, prediction accuracy on conventional benchmarks does not by itself establish that a model has learned the intended physical evidence. A recurring lesson from machine learning is that models can exploit correlations present in training data, including shortcuts or distribution-specific patterns that may not remain reliable under distribution shift^15,16^. Such concerns are especially important in scientific machine learning, where benchmark design and data artifacts can lead to overoptimistic conclusions about model performance^17^. Similar reliability-oriented evaluations have been emphasized in molecular machine learning beyond familiar chemical space and in biological foundation models tested under zero-shot or simple-baseline comparisons^18–20^. For de novo peptide sequencing, this issue is especially relevant because training spectra represent only a small and biased subset of the full theoretical peptide sequence space. In typical proteomics datasets, amino-acid usage, sequence motifs, enzymatic cleavage patterns, and proteome context are all highly structured. Thus, a model may produce plausible peptide sequences partly guided by learned sequence priors, particularly when fragmentation evidence is incomplete or ambiguous. Existing de novo sequencing benchmarks, however, primarily measure prediction accuracy on common proteomics datasets and do not directly quantify prior-associated behavior.

To address this gap, we introduce the Prior Bias Index (PBI), a general framework for quantifying prior-associated model bias as a normalized shift in model behavior from a null baseline toward a reference condition. We implement this framework as DeNovo-PBI, a benchmark for measuring prior-associated behavior in de novo peptide sequencing models. DeNovo-PBI combines controlled peptide-sequence distribution design, *in silico* spectrum generation, spectral-evidence perturbation, and mutation-group analysis to test whether model predictions remain evidence-driven or shift toward learned sequence priors. In this study, we use DeNovo-PBI to evaluate three forms of prior-associated behavior: sequence-distribution dependence, database-associated amino-acid-pair order preference, and prediction consistency under shared sequence context.

We applied DeNovo-PBI to benchmark deep learning-based de novo sequencing models alongside a non-neural de novo sequencing algorithm using random, natural, and mutated peptide sequences paired with *in silico* or experimentally acquired spectra. Across these controlled assays, deep learning models showed sequence-distribution-dependent performance and scoring behavior beyond direct fragment evidence when local fragment evidence was removed. They also exhibited strong directional amino-acid-pair order preferences under order-ambiguous spectral settings, including cases where the fragment ions distinguishing the two possible N-terminal orders were removed. Together, these results show that modern de novo sequencing models can exhibit measurable sequence-prior-associated behavior, and that DeNovo-PBI provides a quantitative benchmark for evaluating such behavior beyond conventional accuracy-based model comparisons.

## Materials and Methods

### Models and datasets

De novo peptide sequencing predictions were generated using three models: the deep learning-based models Casanovo^6,21^ and InstaNovo^13^, and the non-neural de novo sequencing algorithm Novor^9^. Casanovo (version 5.0.0) was run using the model checkpoint casanovo_v5_0_0.ckpt. Predictions were generated with a precursor mass tolerance of 20 ppm, and beam search was configured to use five beams and return up to five beam candidates per spectrum. The Casanovo configuration used carbamidomethylation of cysteine as a fixed modification. InstaNovo (v1.2.2) was run using the transformer-only model with checkpoint instanovo-v1.2.0.ckpt, with de novo prediction enabled, five-beam decoding, a precursor mass tolerance of 20 ppm, and carbamidomethylation of cysteine allowed. Novor (v1.05.0573) was run through DeNovoGUI^22^ (v1.16.8 for Mac and Linux), with a precursor mass tolerance of 20 ppm and fixed carbamidomethylation of cysteine. Model-reported raw scores were retained for downstream score-distribution analyses; for InstaNovo, these scores corresponded to log-probability outputs.

Two broad categories of data were used in this study: computationally generated peptide sequences paired with *in silico* generated spectra, and real peptide sequences with either *in silico* generated spectra or experimentally acquired spectra. Real peptide sequences and spectra were obtained from the curated 9-species dataset^14^ (balanced version v2, https://doi.org/10.5281/zenodo.13685813). To avoid overlap with Casanovo training data, entries with peptide sequences matching the Casanovo training dataset^23^ (version 2, https://doi.org/10.5281/zenodo.12587317) were removed from the 9-species dataset by sequence matching. In addition, Homo sapiens entries were removed, yielding a real-peptide evaluation set that did not contain peptide sequences present in the Casanovo training data or human-derived entries. The detailed design of the peptide and spectrum datasets is described in the following section.

### *In silico* generation of peptide sequences and spectra

*In silico* datasets were designed to isolate sequence-distribution effects and N-terminal order preference under controlled spectral-evidence conditions. To evaluate model behavior under controlled N-terminal sequence ambiguity, we first defined a set of mass-unique unordered two-amino-acid pairs. Candidate pairs were required to contain two distinct amino acids and to have a residue-mass sum that was not within 0.02 Da of any other one-or-more-residue combination. Pairs containing K or R were excluded from the candidate set to maintain a tryptic peptide design, although K- and R-containing combinations were still considered when assessing mass uniqueness. Pairs affected by mass degeneracy, including I/L ambiguity, were therefore excluded by this uniqueness criterion. This procedure yielded 45 mass-unique unordered two-amino-acid pairs, which were placed as the first two N-terminal residues in *in silico*-generated peptides. This design allowed order preference to be evaluated after removal of fragment ions that distinguish the two possible N-terminal residue orders.

Random peptide sequences were generated using the design [mass-unique two-amino-acid prefix] + [random amino-acid sequence] + [R/K]. For each of the 45 two-amino-acid prefixes, 230 random middle sequences of length 17 were generated, with K and R excluded from the middle sequence, and a randomly selected terminal R or K was appended. This produced 10,350 unique peptide sequences of length 20. For each peptide, *in silico* spectra were generated with precursor and fragment charges set to 1+. Two spectrum settings were generated: a complete b/y-ion ladder and a masked ladder in which the b1 ion and complementary y(n−1) ion were removed. An expanded random-peptide dataset was generated using the same design with 23,000 random middle sequences per two-amino-acid prefix, yielding 1,035,000 unique peptide sequences.

Real-peptide *in silico* spectra were generated from the filtered 9-species real-peptide evaluation set described above. Peptides were selected if their unordered first two N-terminal residues matched one of the 45 mass-unique two-amino-acid pairs. This selection contained 28,442 peptide sequences before removal of sequences overlapping the Casanovo training dataset and human-derived entries, and 25,333 peptide sequences after filtering. As in the random-peptide dataset, *in silico* spectra were generated with precursor and fragment charges set to 1+, under both complete b/y-ion ladder and b1- and y(n−1)-removed spectrum settings.

To generate the mutated-peptide dataset, real peptide sequences of length ≥20 were selected as parent peptides for mutation-group construction. Each parent peptide defined one mutation group, in which the N-terminal sequence context was retained across mutated variants derived from the same original peptide. This selection contained 4,341 unique parent peptides before removal of sequences overlapping the Casanovo training dataset and human-derived entries, and 3,890 unique parent peptides after filtering. For each retained parent peptide, 100 mutated variants were generated by replacing 50% of the mutable region from the C-terminal side while preserving the final C-terminal residue. The mutable region was defined as the peptide sequence excluding the final C-terminal amino acid. Replacement residues were randomly sampled with K and R excluded, and the replacement segment was required to differ from the original segment. This design preserved shared N-terminal sequence context within each mutation group while introducing sequence diversity in the C-terminal portion. *In silico* spectra were generated for the mutated peptides using the same precursor and fragment charge settings and the same complete and b1-and y(n−1)-removed spectrum settings described above.

### Model prediction and performance evaluation

Top-1 sequence accuracy was used as the primary performance metric. For complete *in silico* spectra, a prediction was considered correct only if the Top-1 predicted peptide sequence exactly matched the reference sequence. For *in silico* spectra with the b1 ion and complementary y(n−1) ion removed, the first two N-terminal residues were treated as order-ambiguous; a prediction was considered correct if either orientation of the correct first-two-amino-acid composition was followed by an exact match to the remaining reference sequence. For experimentally acquired spectra, accuracy was calculated by comparing the Top-1 predicted sequence with the annotated ground-truth peptide sequence. Prediction-score distributions and amino-acid-level score distributions were analyzed using the original model-reported scores. For beam-level analyses, the top five beam candidates reported by each model were retained when available.

### Prior Bias Index framework

We defined a general Prior Bias Index (PBI) to quantify prior-associated shifts in model behavior under controlled probing conditions. For a bias-probing experiment, let S denote a scalar statistic of model behavior, such as prediction accuracy, prediction frequency, confidence score, or output probability. PBI is defined as

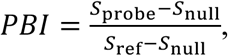

where *S*_probe_ is the statistic measured under the probing condition, *S*_null_ is the statistic measured under a null or prior-minimized baseline, and *S*_ref_ is the statistic measured under a reference condition. PBI measures how far the probing condition shifts away from the null baseline toward the reference condition.

A PBI of 0 indicates null-like behavior, whereas a PBI of 1 indicates that the probing condition reaches the reference level. Positive values indicate movement toward the reference direction, while negative values indicate behavior below or opposite to the null baseline. Values greater than 1 indicate that the probing condition exceeds the reference level. PBI is undefined when *S*_ref_ equals *S*_null_, because no reference difference is available for normalization.

### DeNovo-PBI benchmark

We applied the PBI framework to de novo peptide sequencing and developed the DeNovo-PBI benchmark with three category-specific indices: sequence-distribution PBI, database amino-acid-order PBI, and mutation-group PBI.

Sequence-distribution PBI (PBI_dist_) evaluates whether model performance changes when the peptide sequence distribution changes from random peptides to natural peptides under controlled *in silico* spectral evidence. Let *A*(*D*, *E*) denote the sequence accuracy for peptide distribution *D* under evidence condition *E*. Using the natural peptide distribution *D*_nat_, random peptide distribution *D*_rand_, complete *in silico* spectral evidence *E*_in_silico_, and experimental spectra *E*_exp_, PBI_dist_ was defined as

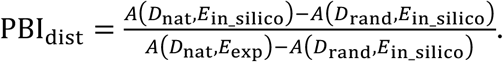

Database amino-acid-order PBI (PBI_DB−order_) evaluates whether model predictions favor amino-acid orders that are more common in the peptide database after order-diagnostic fragment ions are removed. For each unordered N-terminal amino-acid pair {*a*, *b*}, the database-favored order *o*^∗^ was defined as the more frequent order between the two possible adjacent residue orders in the reference database. PBI_DB−order_ was defined as

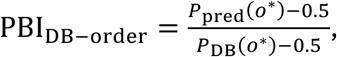

where *P*_pred_(*o*^∗^) is the conditional frequency of predicting the database-favored order among predictions that recovered either orientation of the correct first-two-amino-acid composition, and *P*_DB_(*o*^∗^) is the corresponding conditional database frequency within that unordered pair.

Mutation-group PBI (PBI_mut_) evaluates whether mutated peptides derived from the same parent peptide retain parent-associated sequence patterns despite sequence variation elsewhere in the peptide. In this implementation, the shared pattern examined was parent-consistent N-terminal first-two-amino-acid order after removal of the order-diagnostic fragment ions, namely the b1 ion and complementary y(n−1) ion. For mutation group i, *S_i_* was defined as the fraction of predictions choosing the parent-consistent order among predictions that recovered either orientation of the correct first-two-amino-acid composition under this order-ambiguous spectral setting. PBI_mut_ was defined as

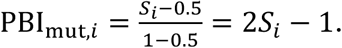

### Statistical analysis

Order-preference significance was assessed using two-sided exact binomial tests against a 50:50 null expectation. For database amino-acid-order analyses, tests were performed for each unordered non-identical amino-acid pair using the number of predictions choosing the database-favored order among predictions that recovered either orientation of the correct first-two-amino-acid composition. For mutation-group analyses, tests were performed for each mutation group using the number of predictions choosing the parent-consistent order among predictions that recovered either orientation of the correct first-two-amino-acid composition.

P-values were adjusted within each experiment, model, condition, and mutation setting using the Benjamini-Hochberg false discovery rate procedure, with significance defined at α = 0.05. Same-residue pairs were excluded from order-bias analyses because the two possible orders are identical. Mutation groups were included in the PBI_mut_ significance analysis only when at least 10 predictions recovered either orientation of the correct first-two-amino-acid composition.

## Results

### DeNovo-PBI enables benchmarking of learned prior bias in de novo peptide sequencing models

Deep learning models for de novo peptide sequencing may learn different sources of predictive information during training, including fragmentation-related spectral patterns and peptide-sequence priors. To benchmark learned sequence priors, we developed DeNovo-PBI (Figure 1), a benchmarking toolkit for evaluating learned prior bias across models. DeNovo-PBI combines benchmark dataset construction, *in silico* sequence and spectral perturbation workflows, PBI-based metrics, and analysis algorithms to compare model behavior across different peptide sequence distributions and evidence settings.

**Figure 1.**
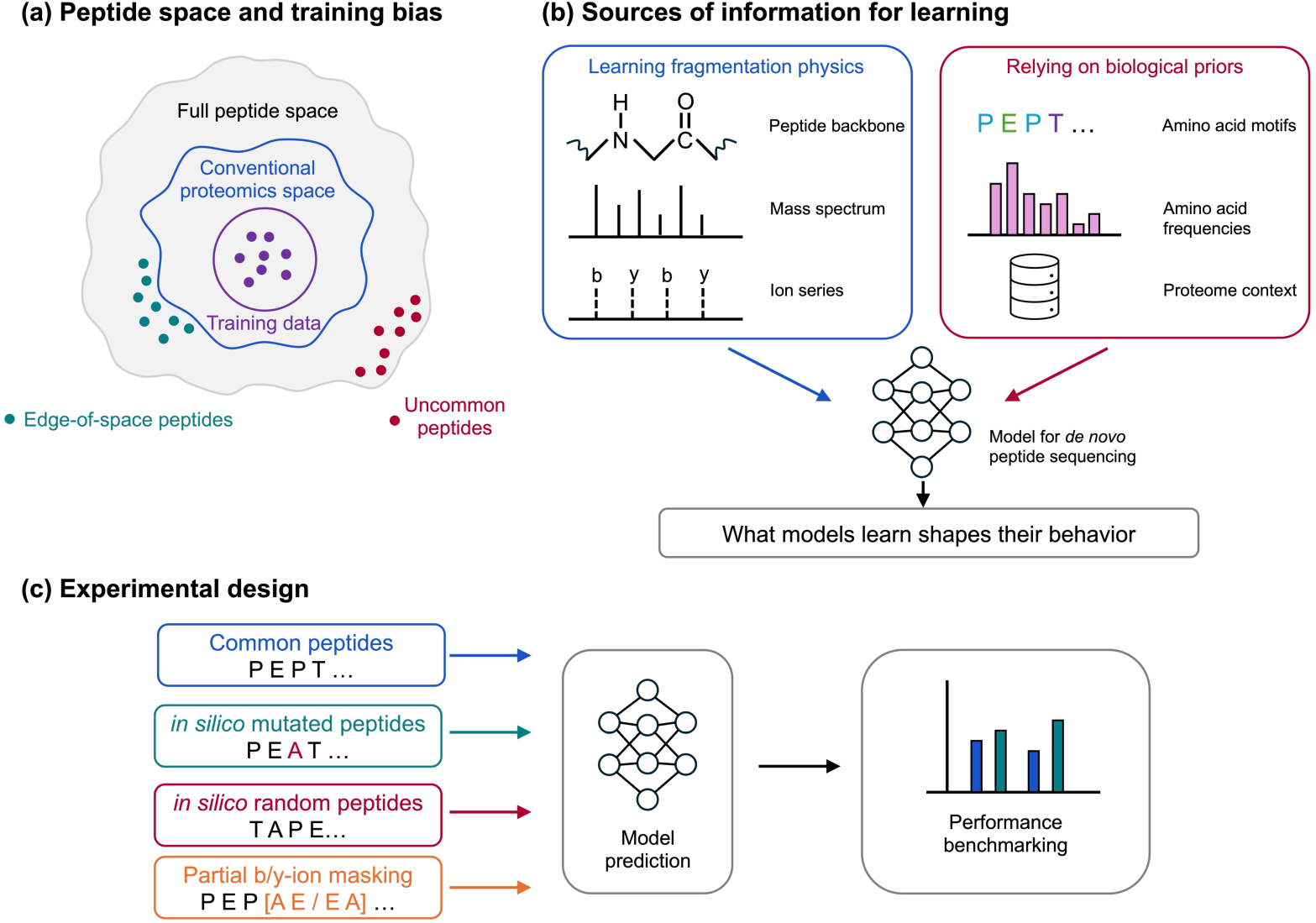
Overview of the DeNovo-PBI framework for benchmarking prior bias in de novo peptide sequencing models. **(a)** Peptide training data occupy only a small and biased subset of the full peptide sequence space, mainly concentrated within conventional proteomics space. Peptides near the boundary of this distribution, or outside conventional proteomic peptide space, provide opportunities to evaluate model behavior beyond the training distribution. **(b)** Deep-learning models for de novo peptide sequencing can learn from multiple sources of information. Models may learn patterns associated with spectral evidence, as well as biological and statistical regularities present in the training data, including amino-acid motifs, residue-frequency patterns, and proteome context. These distinct sources of learnable information motivate the central idea that what models learn shapes how they behave. **(c)** DeNovo-PBI evaluates model behavior across peptide classes with different relationships to the training distribution, including common natural peptides, *in silico* mutated peptides, and *in silico* random peptides. Partial b/y-ion masking further probes order-preference behavior by removing fragment ions that distinguish composition-matched, order-swapped adjacent residue pairs. Differences in model performance and prediction patterns across these controlled evaluations provide a basis for quantifying and benchmarking learned prior bias.

DeNovo-PBI benchmarks model behavior across different sequence distributions to evaluate how well models generalize to unseen peptide sequences. To benchmark model behavior when direct physics-based spectral evidence is absent or ambiguous, we further introduced spectral ambiguity by removing order-diagnostic b/y ions for selected N-terminal amino-acid pairs. These controlled settings allow model behavior to be examined beyond standard real-peptide benchmarks and provide a more complete analysis of model performance.

### Deep learning models show sequence-distribution dependence in peptide prediction

Using the DeNovo-PBI benchmark, we first evaluated whether peptide sequence distribution affected model performance under controlled *in silico* spectral evidence (Figure 2). With complete b/y-ion spectra, the deep learning model Casanovo showed a substantial accuracy increase when the peptide sequence distribution changed from random peptides to real peptides, with Top-1 sequence prediction accuracy increasing from 0.255 to 0.574. This result indicates that Casanovo performance depends strongly on whether peptide sequences follow a natural peptide distribution, even when spectra are generated under controlled *in silico* conditions.

**Figure 2.**
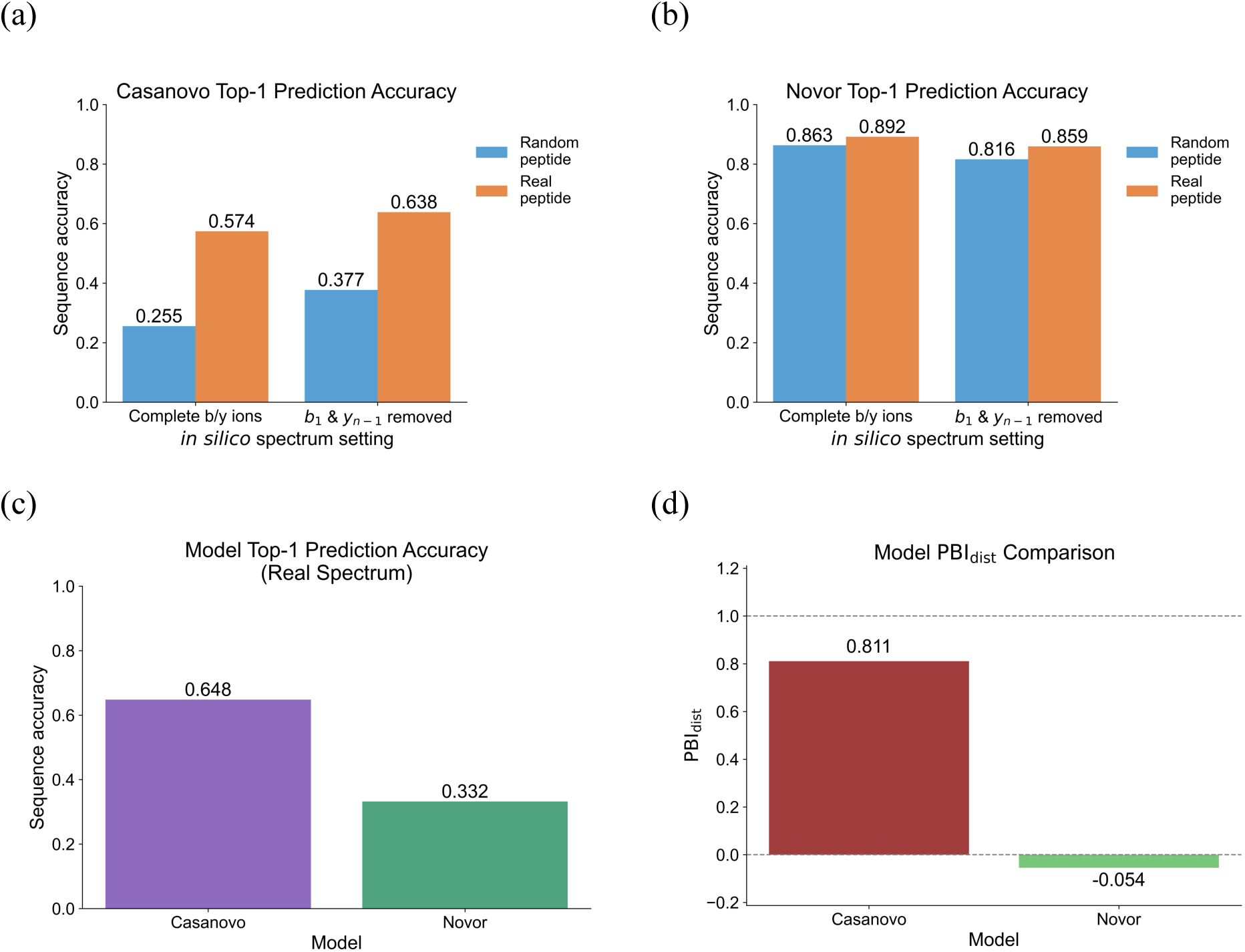
Sequence-distribution prior bias in de novo peptide sequencing models. Top-1 sequence accuracy was evaluated for Casanovo and Novor using *in silico* spectra generated from random peptides and real peptide sequences under controlled fragment-ion settings, together with experimental spectra from real peptides. **(a)** Casanovo Top-1 sequence accuracy on *in silico* spectra generated from random peptides and real peptides with complete b/y ions or with terminal b_1_ and y_n−1_ ions removed. The random-peptide *in silico* dataset contained 10,350 unique spectra for each spectrum setting, and the real-peptide *in silico* dataset contained 25,333 unique spectra. **(b)** Novor Top-1 sequence accuracy on the same *in silico* peptide sequence distributions and spectrum settings. **(c)** Top-1 sequence accuracy of Casanovo and Novor on experimental spectra. Casanovo generated predictions for 25,261 spectra, whereas Novor generated predictions for 25,333 spectra. Values above bars indicate measured Top-1 sequence accuracy. For complete *in silico* spectra, accuracy was defined as an exact sequence match to the reference peptide. For incomplete *in silico* spectra with b_1_ and y_n-1_ ions removed, predictions were considered correct if either ordering of the first two amino acids, followed by an exact match of the remaining sequence, matched the reference. For experimental spectra, accuracy was calculated based on matches to the ground-truth annotations. **(d)** Sequence-distribution Prior Bias Index, PBI_dist_, for each model. PBI_dist_ quantifies how much of the performance shift from the random-peptide *in silico* baseline toward the experimental real-spectrum reference is already observed when the peptide sequence distribution is changed from random to real peptides under complete *in silico* spectral evidence.

Novor, a non-neural-network algorithm, showed a different pattern. Under the same complete b/y-ion setting, Novor achieved high prediction accuracy on both random peptides and real peptides, with comparable accuracies of 0.863 and 0.892. However, on experimentally acquired spectra, Casanovo outperformed Novor, with prediction accuracies of 0.648 and 0.332, respectively. These results show that performance on conventional real-peptide benchmarks does not necessarily predict behavior under shifted peptide sequence distributions, and that models differ in how peptide sequence distribution affects prediction behavior.

We quantified this sequence-distribution dependence using PBI_dist_. Casanovo showed a high PBI_dist_ value of 0.811, whereas Novor showed a near-zero value of −0.054. These results indicate that the performance shift from random peptides toward real-peptide experimental performance was largely captured by changing the sequence distribution for Casanovo, but not for Novor.

Additional analyses supported this pattern. Model-reported Top-1 prediction scores also shifted across peptide sequence distributions and spectrum settings, indicating that sequence distribution affected not only prediction accuracy but also model scoring behavior (Supplementary Figure 1). A length-controlled analysis restricted to peptides of length 20 showed similar model-dependent sequence-distribution effects (Supplementary Figure 2). An expanded random-peptide dataset containing 1,035,000 sequences further confirmed that Casanovo performance on random peptides was consistent at larger scale and was not driven by the smaller random-peptide benchmark size (Supplementary Figure 3). InstaNovo showed behavior comparable to Casanovo, with higher accuracy on real-peptide *in silico* spectra than on random-peptide *in silico* spectra, supporting sequence-distribution dependence across deep learning-based models (Supplementary Figure 4).

### Individual amino-acid scores reveal model reliance on information beyond direct fragment evidence

We next examined whether model-reported amino-acid scores reflected changes in local spectral evidence (Figure 3). For random-peptide *in silico* spectra with complete b/y ions, Casanovo assigned generally high scores across the first five N-terminal positions, especially for the first amino acid. After removal of the b1 ion and complementary y(n−1) ion, Casanovo scores at the N-terminal positions remained high overall, despite the loss of direct order-diagnostic evidence for the first two amino acids.

**Figure 3.**
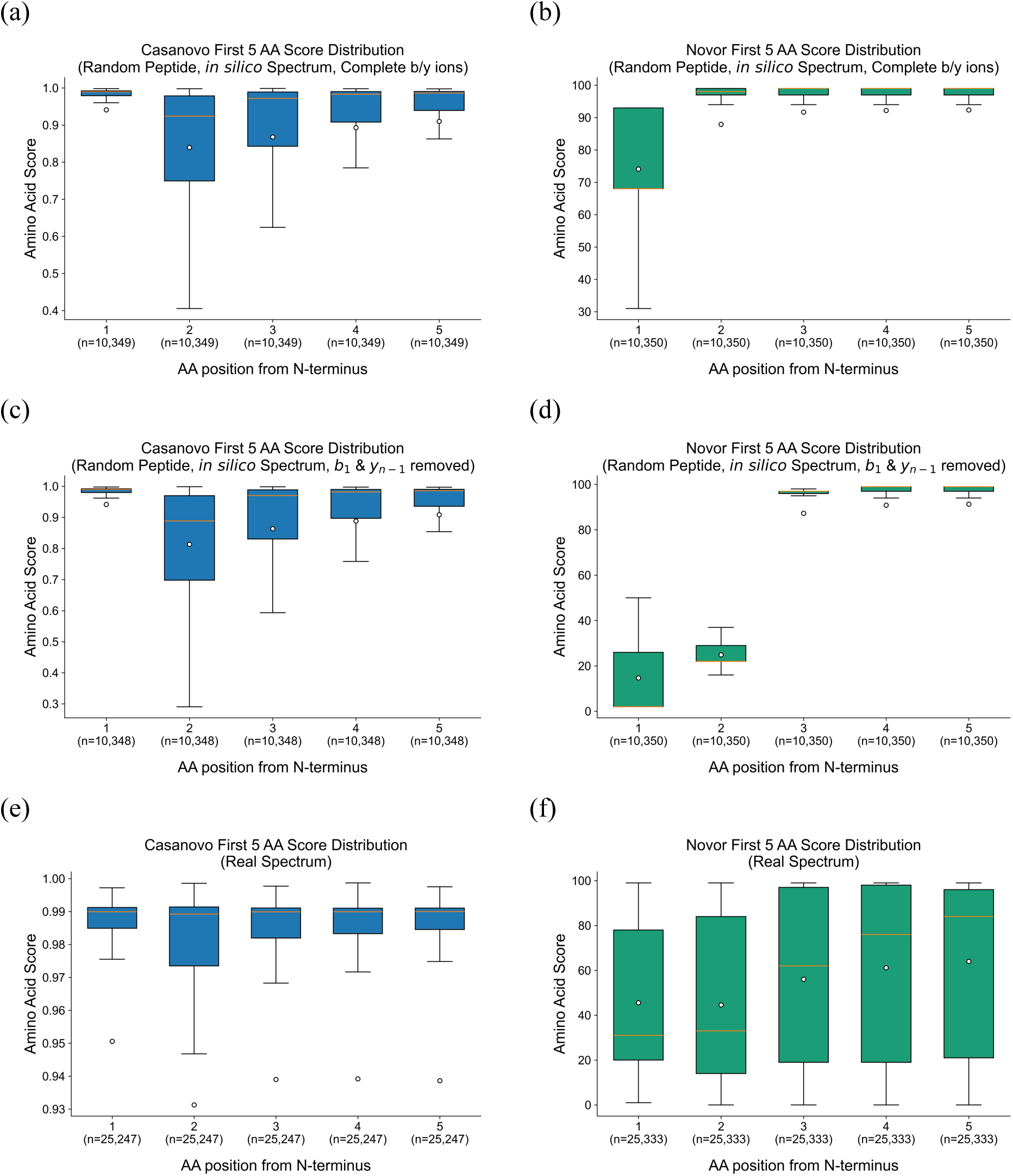
N-terminal amino-acid score distributions across models and spectrum settings. Amino-acid-level score distributions were evaluated for the first five N-terminal positions of Top-1 peptide predictions from Casanovo and Novor. For each model and spectrum setting, only Top-1 predicted sequences of length ≥5 amino acids were included. Scores correspond to the original model-reported amino-acid scores and therefore remain on model-specific native scales. **(a, b)** Score distributions for Casanovo and Novor, respectively, using random-peptide *in silico* spectra with complete b/y ions. **(c, d)** Score distributions for Casanovo and Novor, respectively, using random-peptide *in silico* spectra after removal of terminal b_1_ and y_n-1_ ions. **(e, f)** Score distributions for Casanovo and Novor, respectively, using experimental spectra from real peptides without ion masking. Boxplots summarize the distribution of amino-acid scores at each N-terminal position, and sample sizes below each box indicate the number of Top-1 predictions included for that position.

Novor showed a more evidence-dependent scoring pattern. With complete b/y-ion spectra, Novor assigned high scores to most N-terminal positions, although with greater variability at the first residue. After b1 and complementary y(n−1) ion removal, scores for the first two residues dropped substantially, whereas scores for downstream positions remained high. This pattern is consistent with reduced confidence specifically at positions where direct fragment evidence was removed.

On experimentally acquired spectra, score distributions differed further between models. Casanovo maintained high amino-acid scores across N-terminal positions, whereas Novor showed broad score distributions across the first five residues. This indicates that deep learning models may use spectral information beyond direct b- and y-ion ladder evidence.

Similar patterns were observed on real peptide *in silico* spectra for Casanovo and Novor (Supplementary Figure 5). InstaNovo also showed patterns similar to Casanovo on random-peptide and real-peptide *in silico* spectra (Supplementary Figure 6).

Together, these results show that individual amino-acid scores capture model-specific responses to missing or ambiguous fragment evidence, suggesting that deep learning-based models can rely on learned information beyond direct b- and y-ion series and may assign potentially overconfident predictions even when direct ion-supporting evidence is absent.

### Removal of spectral order-diagnostic ions exposes directional amino-acid order preference

We next tested whether models showed directional order preference when direct spectral evidence distinguishing two N-terminal amino-acid orders was removed (Figure 4). Using random-peptide *in silico* spectra with the b1 ion and complementary y(n−1) ion removed, Casanovo was not expected to distinguish the reference and reversed first-two-amino-acid orders based on direct fragment evidence alone. However, Top-1 predictions showed non-random order behavior: 30.5% matched the reference order recorded in the MGF file, which was alphabetical in this random-peptide setting, 45.0% matched the reversed order, and 24.5% predicted another first-two-amino-acid pair.

**Figure 4.**
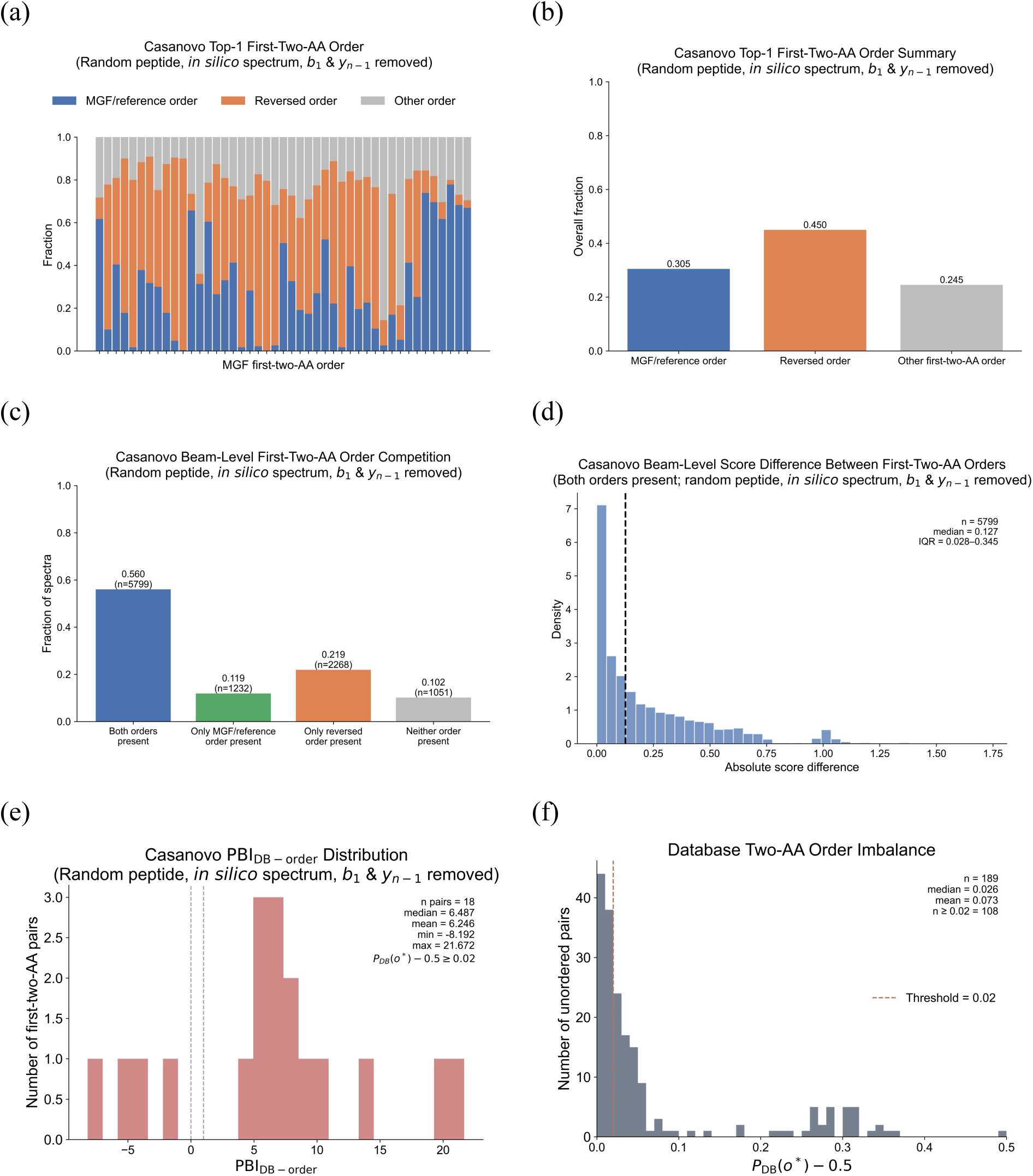
N-terminal two-amino-acid order bias in model predictions after removal of order-diagnostic fragment ions. Order preference in Casanovo predictions was evaluated using 10,350 unique random-peptide *in silico* spectra after removing fragment ions that distinguish the order of the first two N-terminal amino acids. **(a)** Top-1 predicted first-two-amino-acid order across individual N-terminal residue-pair categories. Predictions were classified as matching the MGF/reference order, matching the reversed order, or containing another first-two-amino-acid pair. Fractions are shown within each residue-pair category. **(b)** Overall Top-1 prediction fractions across all evaluated first-two-amino-acid pairs. **(c)** Beam-level competition between the MGF/reference and reversed N-terminal orders. Spectra were grouped according to whether both orders, only the MGF/reference order, only the reversed order, or neither order appeared among the top five beam candidates. **(d)** Distribution of the absolute score difference between MGF/reference-order and reversed-order candidates among spectra for which both orders were present in the top five beam candidates. The dashed line indicates the median score difference, and IQR denotes the interquartile range. **(e)** Distribution of database-order Prior Bias Index (PBI_DB-order_), across significantly order-biased unordered amino-acid pairs. For each unordered pair {a,b}, the database-favored order o* was defined as the more frequent order between ab and ba. PBI_DB-order_ was calculated as PBI_DB-order_ = (*P_pred_*(*o* ∗) − 0.5)/(*P_DB_*(*o* ∗) − 0.5), where *P_pred_*(*o* ∗) denotes the frequency of predicting the database-favored order o* and was computed only among predictions assigning either ab or ba to the first two amino-acid positions. Significance was assessed by two-sided exact binomial tests against 0.5 with Benjamini-Hochberg FDR correction. Among 45 unordered non-identical amino-acid pairs, 39 showed significant deviation from 0.5, and 18 pairs with database order imbalance ≥0.02 are shown. **(f)** Distribution of database order imbalance across unordered non-identical amino-acid pairs. For each pair {a,b}, *P_DB_*(*o* ∗) was defined as the fraction of database occurrences corresponding to the more frequent order between ab and ba, and the imbalance was calculated as *P_DB_*(*o* ∗) − 0.5.

Beam-level analysis further showed that both possible orders were often present among the top five candidates, occurring in 56.0% of spectra. When both orders were present, the absolute score difference was usually small but nonzero, with a median difference of 0.127. In 33.8% of spectra, only one of the two possible orders appeared among the beam candidates. These results indicate that Casanovo showed directional preference when spectral evidence was ambiguous.

We quantified this behavior using database amino-acid-order PBI (PBI_DB−order_), using the Casanovo training dataset as the reference database for defining database-favored amino-acid orders. Among 45 unordered non-identical amino-acid pairs, 39 showed significant deviation from the 0.5 null expectation for prediction frequency, indicating that the two possible orders were not predicted equally often after order-diagnostic ions were removed. Among significantly order-biased amino-acid pairs with database order imbalance ≥0.02, PBI_DB−order_ values were largely positive, with a median of 6.487, indicating that predictions often shifted toward the database-favored order even after removal of direct order-diagnostic ions. Similar analyses using real-peptide *in silico* spectra and InstaNovo supported the presence of model-dependent order behavior under order-ambiguous spectral settings (Supplementary Figures 7 and 8).

### Mutated peptide groups reveal consistent order preference under shared sequence context

We next tested whether model predictions remained consistent within groups of mutated peptides derived from the same parent sequence (Figure 5). In this design, mutated peptides shared the same N-terminal sequence context but contained sequence variation elsewhere in the peptide. After removal of the b1 ion and complementary y(n−1) ion, the order of the first two N-terminal amino acids was no longer directly resolved by fragment evidence.

**Figure 5.**
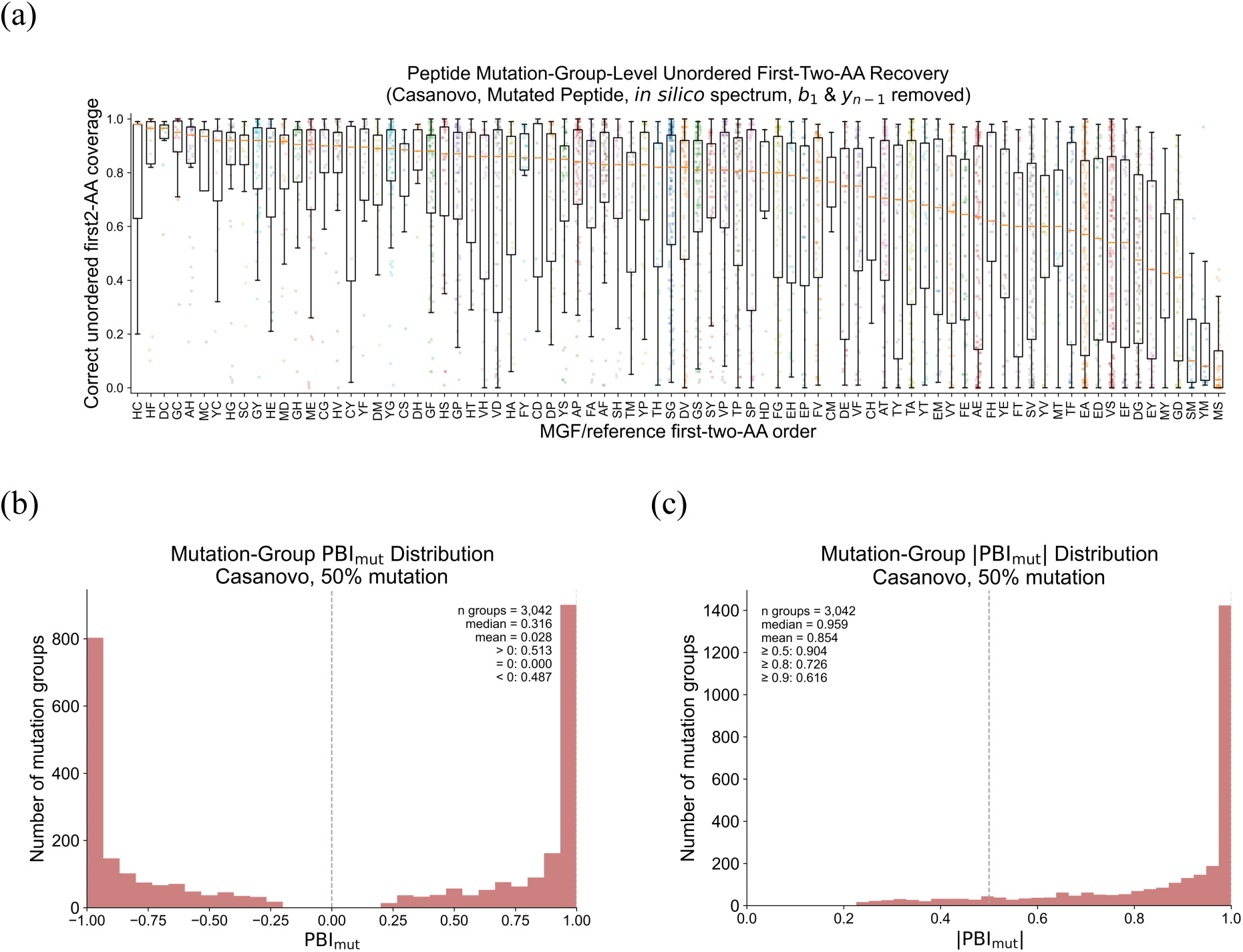
Mutation-group-level N-terminal two-amino-acid order bias in Casanovo predictions for mutated peptides with shared sequences. Casanovo predictions were evaluated using *in silico* spectra generated from mutated peptide sequences derived from real peptides. Mutated peptides were generated from peptides of length ≥20 by mutating 50% of the mutable region from the C-terminal side while preserving the final C-terminal residue. Each original peptide sequence (backbone) was designed to produce 100 mutated variants. **(a)** Mutation-group-level recovery of the correct unordered N-terminal first-two-amino-acid composition across 3,890 parent-peptide backbones. Each point represents one mutation group, and the y-axis indicates the fraction of retained mutated spectra for which the Top-1 prediction matched either orientation of the parent first-two-amino-acid pair. Boxplots summarize mutation groups within each parent first-two-amino-acid order. **(b)** Distribution of mutation-group Prior Bias Index (PBI_mut_) among significantly order-biased mutation groups. PBI_mut_quantifies whether mutated peptides derived from the same parent sequence retain the parent-consistent N-terminal first-two-amino-acid order. For mutation group *i*, PBI_mut,i_ = 2*S_i_* − 1, where *S_i_* is the fraction of predictions choosing the parent-consistent order among predictions that recovered either orientation of the correct first-two-amino-acid composition. Of 3,433 groups with at least 10 predictions that recovered either orientation of the correct first-two-amino-acid composition, 3,042 showed significant deviation from the 50:50 null by two-sided exact binomial testing with Benjamini-Hochberg FDR correction at α = 0.05 and are shown. **(c)** Distribution of |PBI_mut_| for the same groups.

Across 3,890 parent-peptide-derived mutation groups, Casanovo often recovered the correct unordered first-two-amino-acid composition, but the recovery rate varied across parent N-terminal amino-acid pairs. We then used PBI_mut_ to quantify whether predictions within each mutation group consistently favored one of the two possible N-terminal orders. Among 3,433 eligible mutation groups with sufficient predictions recovering the correct unordered first-two-amino-acid composition, 3,042 showed significant deviation from the 50:50 null expectation.

The distribution of PBI_mut_showed strong group-level order preference, but not exclusively in the parent-consistent direction. These results indicate that, under order-ambiguous spectral evidence, Casanovo predictions can follow consistent shared-sequence-associated behavior within mutated peptide groups, even when the direction of the preferred order differs across groups.

## Discussion

In this study, we developed DeNovo-PBI to quantify prior-associated behavior in de novo peptide sequencing models under controlled sequence and spectral perturbations. Our results show that deep learning-based de novo sequencing models can exhibit measurable dependence on peptide sequence priors, in addition to their use of fragmentation-related spectral evidence. This observation raises a fundamental benchmarking question for modern de novo sequencing: to what extent are predicted peptide sequences inferred from the acquired spectrum itself, rather than shaped by sequence regularities and dataset-specific biases learned from training data? Conventional accuracy-centered benchmarks cannot fully answer this question, because they evaluate whether predictions match known peptide labels but do not directly test how model behavior changes when spectral evidence is incomplete, ambiguous, or distributionally shifted. DeNovo-PBI therefore provides an essential benchmark for evaluating prior-associated behavior in de novo peptide sequencing.

The strongest evidence for prior-associated behavior came from assays in which peptide sequence distribution or local spectral evidence was systematically controlled. Casanovo and InstaNovo performed substantially better on natural peptide sequences than on random peptide sequences under matched *in silico* spectral settings, whereas Novor showed much less sequence-distribution dependence. When fragment ions that distinguish the two possible N-terminal amino-acid orders were removed, deep learning models still showed directional order preferences, often shifting toward database-favored residue orders. Mutation-group analyses further showed that predictions could remain consistently biased within groups sharing the same sequence context, despite sequence variation elsewhere in the peptide. Together, these results indicate that learned peptide-sequence priors can measurably influence de novo sequencing behavior, especially when spectral evidence is incomplete or ambiguous.

These findings have important implications for applications that rely on de novo sequencing to explore peptide sequences outside well-characterized databases. In settings such as variant peptide discovery, immunopeptidomics, non-model organism proteomics, and modified peptide analysis, the target peptide distribution may differ substantially from the proteomics datasets used to train current models. In such cases, learned sequence priors may improve predictions when they reflect true biological regularities, but they may also bias predictions toward familiar peptide patterns when the correct sequence is rare, unexpected, or weakly supported by fragment evidence. Prior-aware benchmarking should therefore complement standard peptide- and amino-acid-level accuracy evaluation, especially when de novo predictions are used as evidence for novel biological sequences.

This study also has limitations that define opportunities for extending DeNovo-PBI. The database-associated order analysis uses amino-acid-pair frequencies as a reference prior, but pair-order frequency differences in the database are often small; therefore, the measured order preference likely reflects not only local pair frequencies, but also broader sequence context and other regularities learned during training. The mutation-group assay further showed strong within-group order preference under shared sequence context, although this preference was not systematically aligned with the parent N-terminal order, suggesting that the assay captures context-dependent consistency rather than a single global directional bias. Future designs could examine longer conserved sequence contexts and perturb diagnostic ions at additional peptide positions beyond the N terminus. In addition, our *in silico* spectra were intentionally simplified using uniform fragment-ion intensities to isolate sequence-order ambiguity, but future implementations could incorporate predicted fragment intensities, additional charge states, fragmentation types, and peptide modifications. More broadly, the assays presented here should be viewed as controlled examples of the DeNovo-PBI framework, which can be extended to probe additional forms of learned sequence priors and to support increasingly realistic robustness benchmarks for de novo peptide sequencing models.

More generally, PBI offers a flexible strategy for evaluating whether model behavior shifts toward learned priors under controlled perturbations. Although DeNovo-PBI focuses on de novo peptide sequencing, the same principle could be extended to other deep learning tasks where predictions reflect both input evidence and training-distribution regularities. By pairing task-specific perturbations with appropriate null and reference conditions, PBI-based benchmarks can help evaluate learned bias across broader scientific machine learning applications. Such evaluations may support the development of more robust, interpretable, and evidence-grounded models.

## Supplementary Figures

**Supplementary Figure 1.**
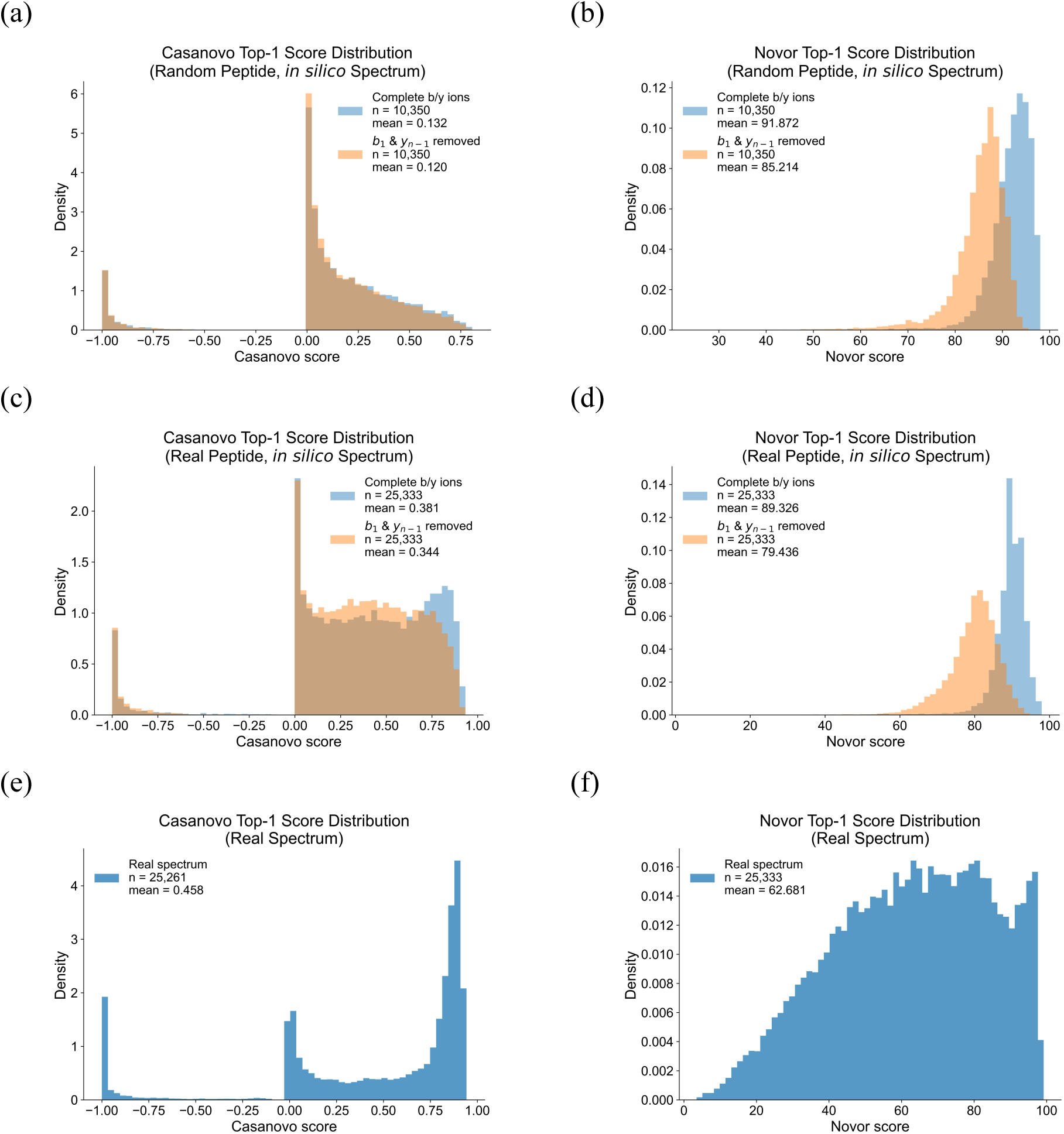
Top-1 prediction score distributions across models, peptide sequence distributions, and spectrum settings. Top-1 prediction scores were evaluated for Casanovo and Novor using the original model-reported raw scores, which remain on model-specific native scales. Histograms show density distributions of Top-1 scores under different peptide and spectrum settings. **(a, b)** Score distributions for Casanovo and Novor, respectively, using random-peptide *in silico* spectra with complete b/y ions or with b_1_ and y_n-1_ ions removed. **(c, d)** Score distributions for Casanovo and Novor, respectively, using real-peptide *in silico* spectra with complete b/y ions or with b_1_ and y_n-1_ ions removed. **(e, f)** Score distributions for Casanovo and Novor, respectively, using real spectra.

**Supplementary Figure 2.**
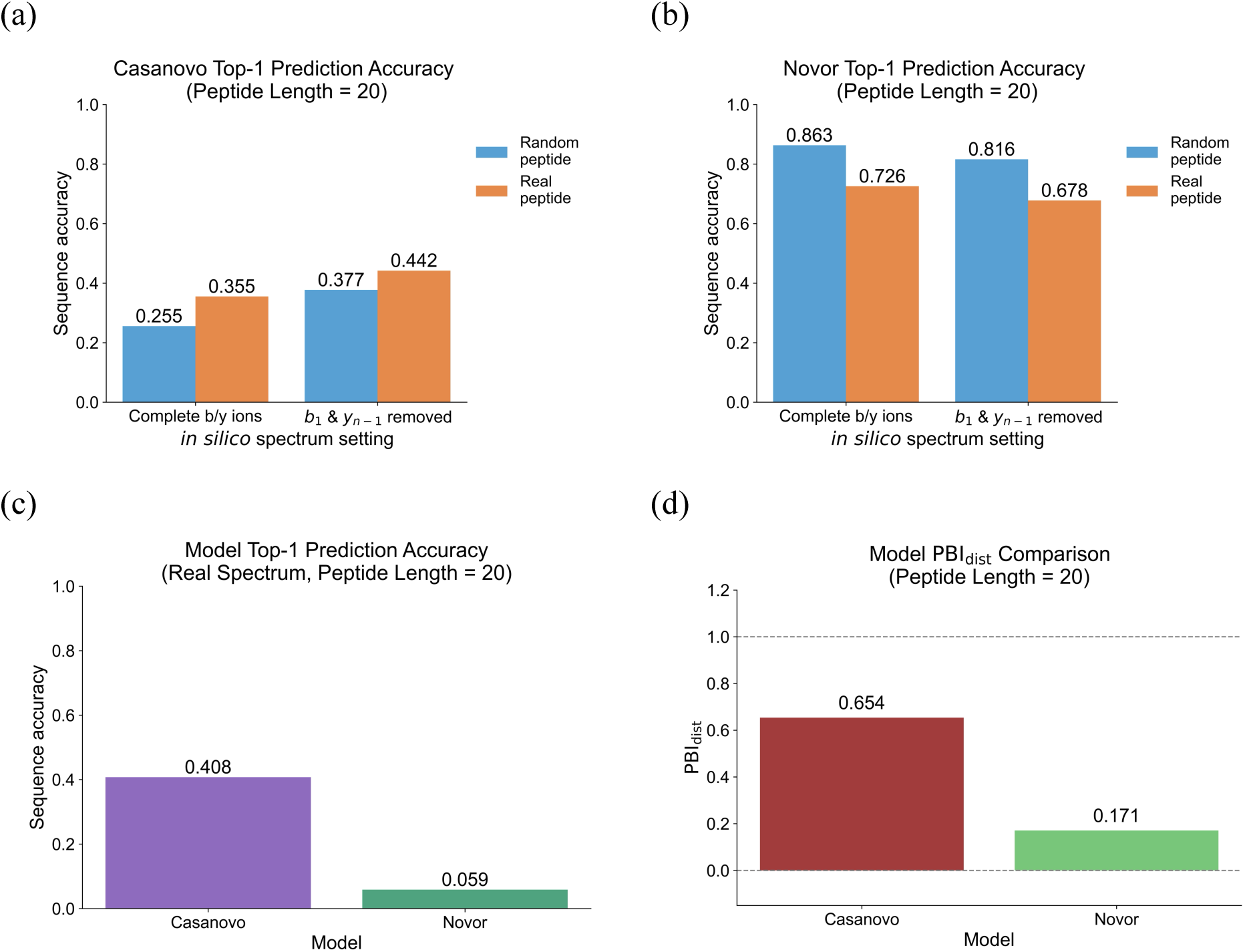
Length-controlled evaluation of Top-1 prediction accuracy and sequence-distribution prior bias for peptides of length 20. Top-1 sequence accuracy was evaluated after restricting datasets to peptides with sequence length equal to 20. Random-peptide *in silico* spectra were generated with peptide length 20, whereas the real-peptide *in silico* and experimental-spectrum datasets each contained 645 peptide-spectrum matches. **(a)** Casanovo Top-1 prediction accuracy on *in silico* spectra from random peptides and real peptides with complete b/y ions or with b_1_ and y_n-1_ ions removed. **(b)** Novor Top-1 prediction accuracy on the same length-restricted *in silico* datasets and spectrum settings. **(c)** Top-1 prediction accuracy of Casanovo and Novor on experimental spectra from peptides with length 20. Values above bars indicate measured Top-1 sequence accuracy. **(d)** PBI_dist_ comparison under the peptide-length-controlled setting.

**Supplementary Figure 3.**
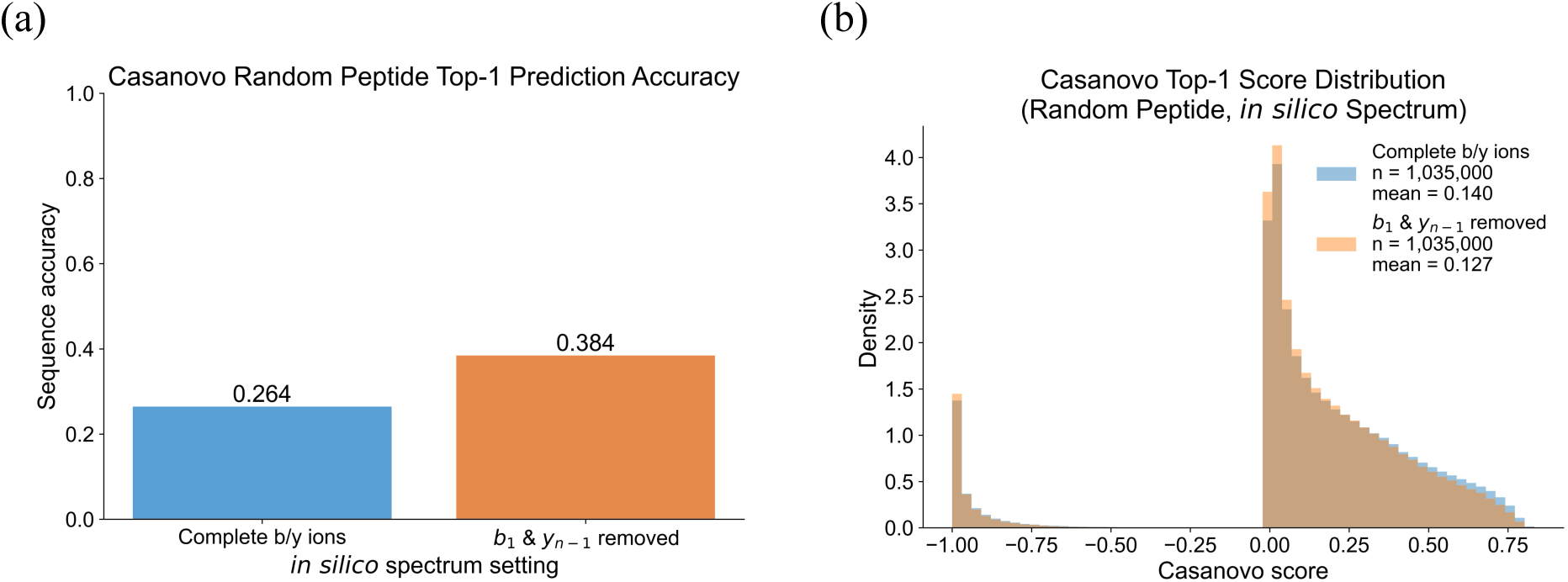
Casanovo performance and score distributions on an expanded random-peptide *in silico* dataset. Casanovo Top-1 predictions were evaluated using 1,035,000 random-peptide *in silico* spectra under two spectrum settings: complete b/y-ion spectra and spectra with the b_1_ ion and complementary y_n−1_ ion removed. **(a)** Top-1 prediction accuracy for random-peptide spectra with complete b/y ions or with b_1_ and y_n-1_ ions removed. Values above bars indicate the measured Top-1 prediction accuracy. **(b)** Distribution of Casanovo Top-1 prediction scores under the same spectrum settings. Scores correspond to the original model-reported raw scores, and sample sizes and mean scores are shown in the plot legend.

**Supplementary Figure 4.**
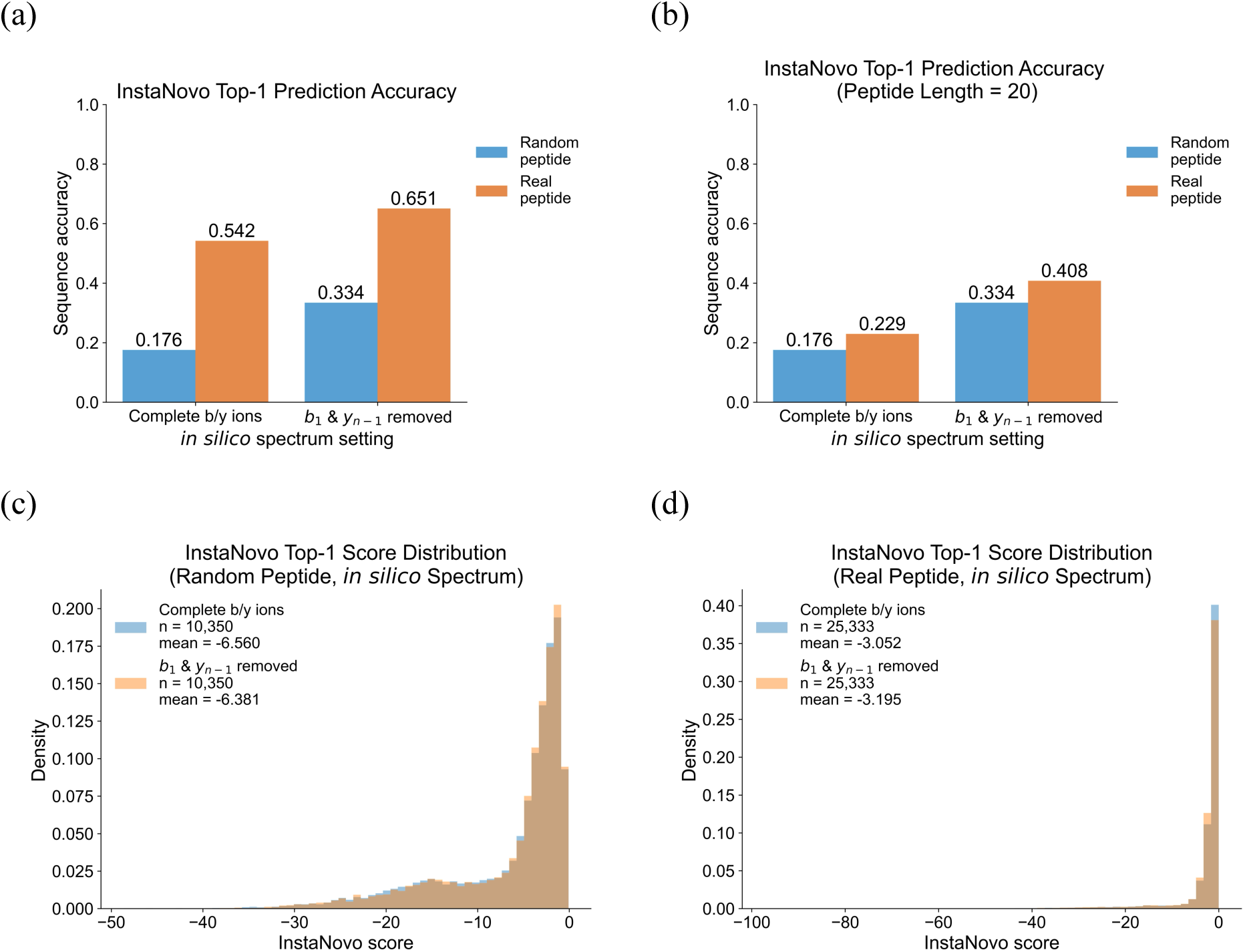
InstaNovo Top-1 prediction accuracy and score distributions across peptide sequence distributions and spectrum settings. InstaNovo Top-1 predictions were evaluated using *in silico* spectra generated from random peptides and real peptide sequences under two spectrum settings: complete b/y-ion spectra and spectra with the b_1_ ion and complementary y_n−1_ ion removed. **(a)** Top-1 prediction accuracy for random-peptide and real-peptide *in silico* spectra across the two spectrum settings. **(b)** Top-1 prediction accuracy after restricting the datasets to peptides with sequence length equal to 20. **(c, d)** Distributions of InstaNovo Top-1 prediction scores (log probabilities) for random-peptide and real-peptide *in silico* spectra, respectively, under two spectrum settings. Values above bars indicate the measured Top-1 prediction accuracy, and sample sizes and mean scores are shown in the plot legends.

**Supplementary Figure 5.**
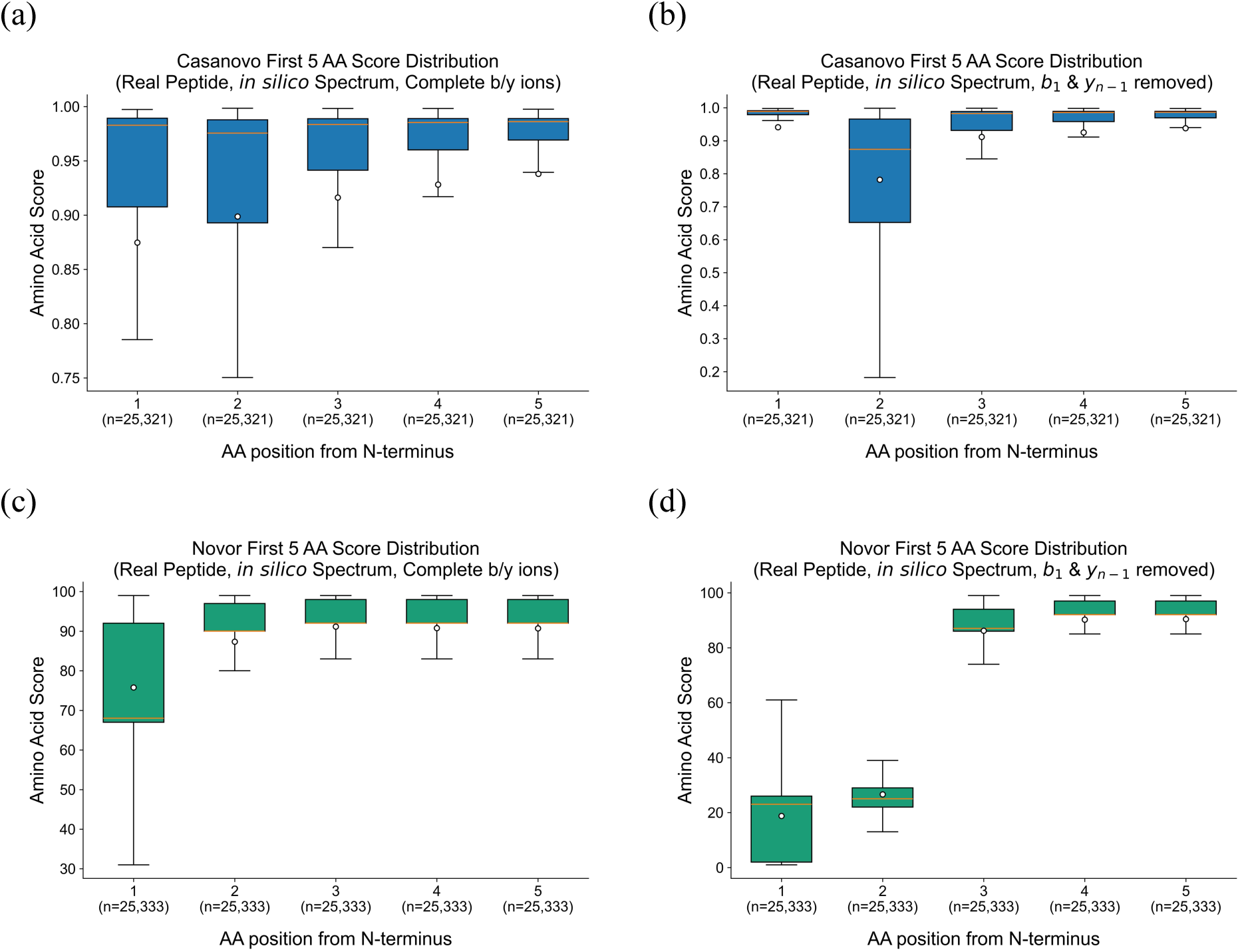
N-terminal amino acid score distributions for real-peptide *in silico* spectra across models and spectrum settings. Amino-acid-level score distributions were evaluated for the first five N-terminal positions of Top-1 peptide predictions from Casanovo and Novor using real-peptide *in silico* spectra. Only Top-1 predicted sequences of length ≥5 amino acids were included. Scores correspond to the original model-reported raw amino acid scores. **(a, b)** Score distributions for Casanovo using complete b/y-ion spectra and spectra with the b_1_ ion and complementary y_n−1_ ion removed, respectively. **(c, d)** Score distributions for Novor using complete b/y-ion spectra and spectra with the b_1_ ion and complementary y_n−1_ ion removed, respectively. Boxplots summarize the distribution of amino acid scores at each N-terminal position; sample sizes below each box indicate the number of Top-1 predictions included for that position.

**Supplementary Figure 6.**
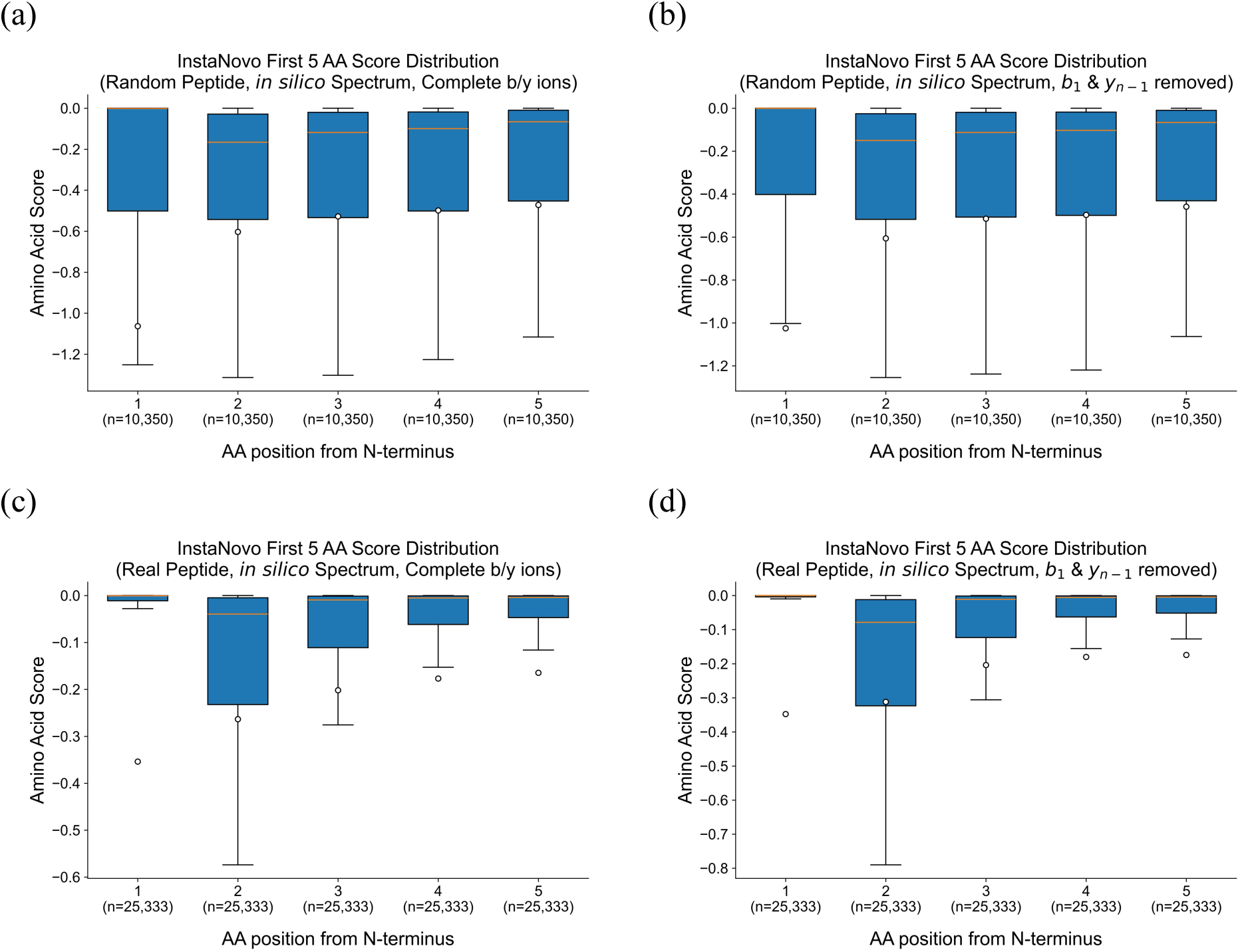
N-terminal amino acid score distributions for InstaNovo Top-1 predictions across peptide sequence distributions and spectrum settings. Amino-acid-level score distributions were evaluated for the first five N-terminal positions of InstaNovo Top-1 peptide predictions. Only Top-1 predicted sequences of length ≥5 amino acids were included. Scores correspond to the original model-reported amino-acid-level scores, reported as log probabilities. **(a, b)** Score distributions for random-peptide *in silico* spectra with complete b/y ions and with the b_1_ ion and complementary y_n−1_ ion removed, respectively. **(c, d)** Score distributions for real-peptide *in silico* spectra with complete b/y ions and with the b_1_ ion and complementary y_n−1_ ion removed, respectively. Boxplots summarize the distribution of amino acid scores at each N-terminal position; sample sizes below each box indicate the number of Top-1 predictions included for that position.

**Supplementary Figure 7.**
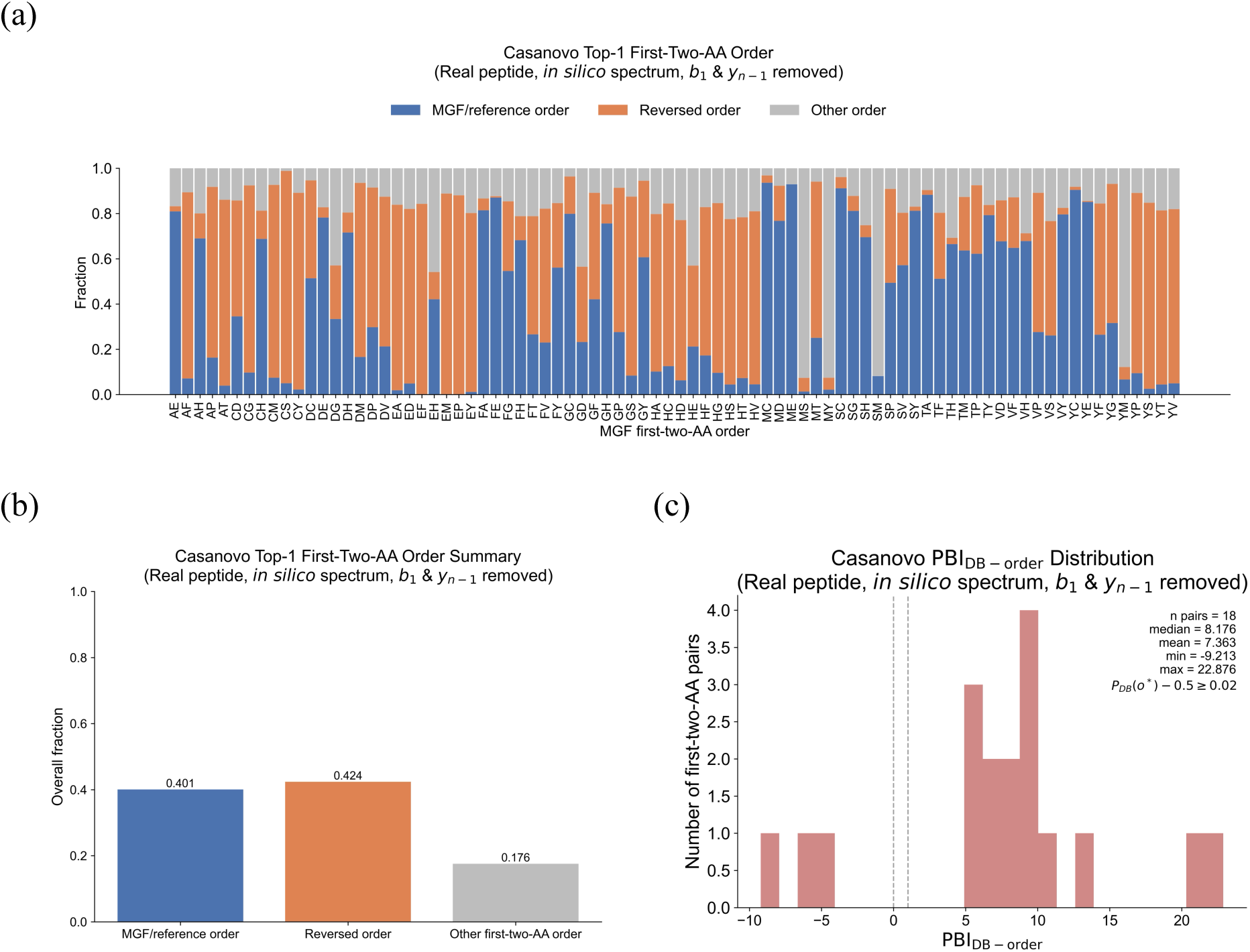
N-terminal two-amino-acid order bias in Casanovo predictions for real-peptide *in silico* spectra. Order preference was evaluated using real-peptide *in silico* spectra after removal of terminal b_1_ and y_n−1_ ions that distinguish the order of the first two N-terminal amino acids. **(a)** Top-1 predicted first-two-amino-acid order across individual N-terminal residue-pair categories. Predictions were classified as matching the MGF/reference order, matching the reversed order, or containing another first-two-amino-acid pair. **(b)** Overall Top-1 prediction fractions across all evaluated first-two-amino-acid pairs. **(c)** Distribution of database-order Prior Bias Index (PBI_DB-order_) across selected unordered amino-acid pairs, computed as in Figure 4.

**Supplementary Figure 8.**
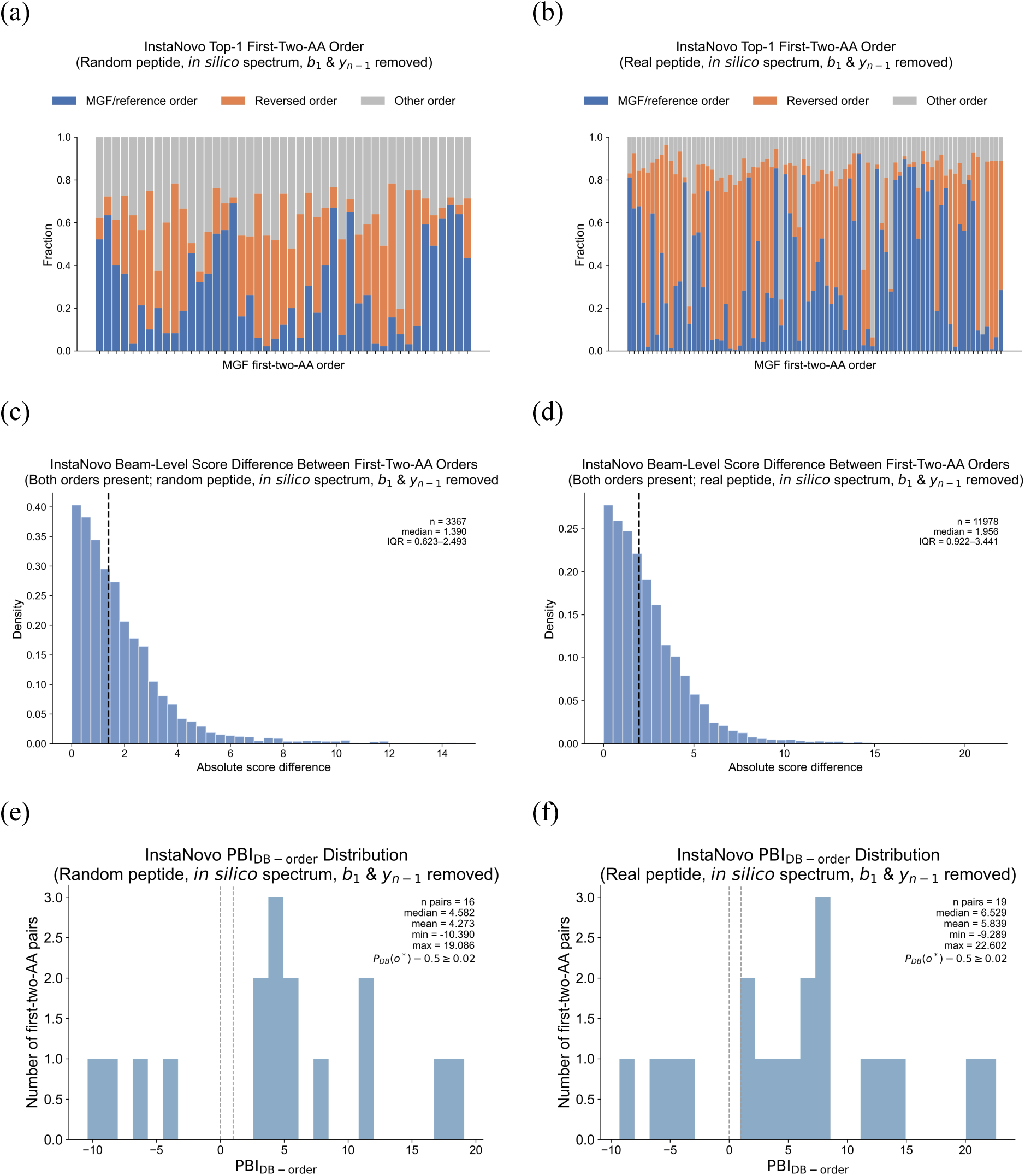
N-terminal first-two-amino-acid order bias in InstaNovo predictions after removal of order-diagnostic fragment ions. InstaNovo predictions were evaluated using random-peptide and real-peptide *in silico* spectra after removal of terminal b_1_ and y_n−1_ ions that distinguish the order of the first two N-terminal amino acids. **(a, b)** Top-1 predicted first-two-amino-acid order across individual residue-pair categories for random-peptide and real-peptide *in silico* spectra, respectively. Predictions were classified as matching the MGF/reference order, matching the reversed order, or containing another first-two-amino-acid pair. **(c, d)** Distribution of the absolute score difference between MGF/reference-order and reversed-order candidates among spectra for which both orders were present in the top five beam candidates, for random-peptide and real-peptide *in silico* spectra, respectively. Score differences were calculated using the original model-reported log-probability scores. The dashed line indicates the median score difference, and IQR denotes the interquartile range. **(e, f)** Distribution of database-order Prior Bias Index (PBI_DB-order_), across significantly order-biased unordered amino-acid pairs, for random-peptide and real-peptide *in silico* spectra, respectively.

